# Bayesian hierarchical modeling of size spectra

**DOI:** 10.1101/2023.02.14.528491

**Authors:** Jeff S. Wesner, Justin P.F. Pomeranz, James R. Junker, Vojsava Gjoni

**Affiliations:** University of South Dakota, Department of Biology, Vermillion, SD 57069; Colorado Mesa University, Department of Physical and Environmental Sciences, Grand Junction, CO 81501; Great Lakes Research Center, Michigan Technological University, Houghton, MI 49931; Louisiana Universities Marine Consortium, Chauvin, LA 70344

**Keywords:** Bayesian, body size spectra, hierarchical, Pareto, power law, Stan

## Abstract

1. A fundamental pattern in ecology is that smaller organisms are more abundant than larger organisms. This pattern is known as the individual size distribution (ISD), which is the frequency of all individual body sizes in an ecosystem.
2. The ISD is described by a power law and a major goal of size spectra analyses is to estimate the exponent of the power law, λ. However, while numerous methods have been developed to do this, they have focused almost exclusively on estimating λ from single samples.
3. Here, we develop an extension of the truncated Pareto distribution within the probabilistic modeling language Stan. We use it to estimate multiple λs simultaneously in a hierarchical modeling approach.
4. The most important result is the ability to examine hypotheses related to size spectra, including the assessment of fixed and random effects, within a single Bayesian generalized (non)-linear mixed model. While the example here uses size spectra, the technique can also be generalized to any data that follows a power law distribution.

## Introduction

In any ecosystem, large individuals are typically more rare than small individuals. This fundamental feature of ecosystems leads to a remarkably common pattern in which relative abundance declines with individual body size, generating the individual size distribution (ISD), also called the community size spectrum (Platt & Denman 1977; Sprules *et al*. 1983; White *et al*. 2008). Understanding how body sizes are distributed has been a focus in ecology for over a century (Sheldon & Parsons 1967; Kerr 1974; Peters & Wassenberg 1983), in part because body sizes distributions reflect fundamental measures of ecosystem structure and function, such as trophic transfer efficiency (Kerr & Dickie 2001; White *et al*. 2007; Perkins *et al*. 2019). Individual size distributions are also predicted as a result of physiological limits associated with body size, thereby emerging from predictions of metabolic theory and energetic equivalence (Brown *et al*. 2004).

More formally, the ISD is a probability density function with a single free parameter λ, corresponding to the following probability density function (Edwards *et al*. 2020):

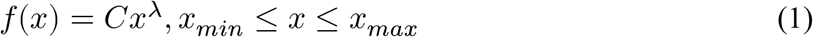

where *x* is the body size (e.g., mass or volume) of an individual regardless of taxon, *x*_*min*_ is the smallest individual attainable and *x*_*max*_ is the largest possible individual (White *et al*. 2008). *C* is a constant equal to:

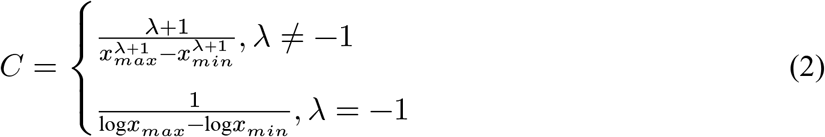

This model is also known as the bounded power law or truncated Pareto distribution. The terms “bounded” or “truncated” refer to the limits of *x*_*min*_ and *x*_*max*_. In practice, values of *x*_*min*_ and *x*_*max*_ often come from the minimum and maximum body sizes in a data set or are estimated statistically (White *et al*. 2008; Edwards *et al*. 2017).

A compelling feature of size spectra is that λ may vary little across ecosystems as a result of physiological constraints that lead to size-abundance patterns more broadly. Metabolic scaling theory predicts 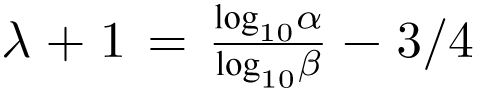, where *α* is trophic transfer efficiency in the food web and *β* is the mean predator-prey mass ratio (Reuman *et al*. 2008). The value of −3/4 is the scaling coefficient of metabolic rate and mass (0.75) (Brown *et al*. 2004) and as a result, values of λ have been used to estimate metabolic scaling across ecosystems (Reuman *et al*. 2008; Perkins *et al*. 2018, 2019). Values of ∼-2 represent a reasonable first guess of expected ISD exponents, with values of ranging from -1.2 to -2 appearing in the literature (Andersen & Beyer 2006; Blanchard *et al*. 2009; Pomeranz *et al*. 2022).

Whether λ represents a fixed or variable value is debated, but it varies among samples and ecosystems (Blanchard *et al*. 2009; Perkins *et al*. 2018; Pomeranz *et al*. 2022). It is often described by its connection with the steepness of log-log plots of size spectra, with more negative values (i.e., “steeper”) indicating lower abundance of large relative to small individuals, and vice versa. These patterns of size frequency are an emergent property of demographic processes (e.g., age-dependent mortality), ecological interactions (e.g., size-structured predation, trophic transfer efficiency), and physiological constraints (e.g., size-dependent metabolic rates) (Muller-Landau *et al*. 2006; Andersen & Beyer 2006; White *et al*. 2008). As a result, variation in λ across ecosystems or across time can indicate fundamental shifts in community structure or ecosystem functioning. For example, overfishing in marine communities has been detected using size spectra in which λ was steeper than expected, indicating fewer large fish than expected (Jennings & Blanchard 2004). Shifts in λ have also been used to document responses to acid mine drainage in streams (Pomeranz *et al*. 2019), land use (Martínez *et al*. 2016), resource subsidies (Perkins *et al*. 2018), and temperature (O’Gorman *et al*. 2017).

Given the ecological information it conveys, the data required to estimate size spectra a vector of individual body sizes are deceptively simple. As long as the body sizes are collected systematically and without bias towards certain sizes, there is no need to know any more ecological information about the data points (e.g., trophic position, age, abundance). However, the statistical models used to estimate λ are diverse. Edwards *et al*. (2017) documented eight methods. Six involved binning, in which the body sizes are grouped into size bins (e.g., 2-50 mg, 50-150 mg, etc.) and then counted, generating values for abundance within each size bin. Binning and log-transformation allows λ to be estimated using simple linear regression. Unfortunately, the binning process also removes most of the variation in the data, collapsing information from 1000’s of individuals into just 6 or so bins. Doing so can lead to inaccurate values of λ, sometimes drastically so (Goldstein *et al*. 2004; White *et al*. 2008; Pomeranz *et al*. 2023).

An improved alternative to binning and linear regression is to fit the body size data to a power law probability distribution (White *et al*. 2008; Edwards *et al*. 2017; Edwards *et al*. 2020). This method uses all raw data observations directly to estimate λ, typically using the maximum likelihood estimation method (Edwards *et al*. 2017). In addition to estimating size spectra of single samples, ecologists have used this method to examine how λ varies across environmental gradients (Perkins *et al*. 2019; Pomeranz *et al*. 2022). However, these analyses typically proceed in two steps. First, λ is estimated individually from each collection (e.g., each site or year, etc.). Second, the estimates are used as response variables in a linear model to examine how they relate to corresponding predictor variables (Edwards *et al*. 2020). We refer to this as the “two-step” approach. A downside to the two-step approach is that it treats body sizes (and subsequent λ’s) as independent samples, even if they come from the same site or time. It also removes information on sample size (number of individuals) used to derive λ. As a result, the approach not only separates the data generation model from the predictor variables, but it is unable to take advantage of partial pooling during model fitting.

Here, we develop a Bayesian modelling framework that uses the truncated Pareto distribution to estimate λ in response to both fixed and random predictor variables. The primary benefit of this approach is that it combines the data generation process and the linear (or non-linear) model into a single generalized linear or non-linear mixed model. The model extends the maximum likelihood approach developed by Edwards *et al*. (2020) to allow for a flexible hierarchical structure, including partial pooling, within the modeling language Stan (Stan Development Team 2022).

## Methods

### Translating to Stan

Stan is a probabilistic modeling language that estimates Bayesian posteriors using Hamiltonion Monte Carlo (Stan Development Team 2022). It does not contain the truncated Pareto described in Eq. 2, so we added it as a user defined log probability density function (lpdf) in rstan Stan Development Team (2022) (Appendix S1). The lpdf was slightly modified as described in S.1.4 of Edwards *et al*. (2020) to contain a term for the count of each body size. For example, if a sample of body sizes is *x* = {1.2, 1.2, 1.5, 2.8}, then these can be re-formatted to include a vector for all unique *x* values {1.2, 1.5, 2.8} and a vector for their *counts* = {2,1,1}, where *counts* can also be non-integers. The benefit of allowing non-integers is that *counts* can reflect densities or catch rates (e.g, no/*m*^2^ or no/hr) in addition to raw counts (Edwards *et al*. 2020).

Converting Eq. 2 into Stan allows for Bayesian estimation of λ’s using generalized (non)-linear mixed models. For example, an intercept-only model would look like this:

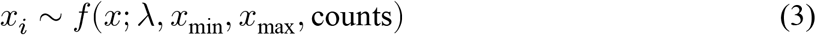

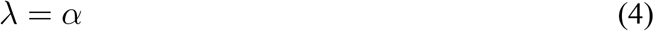

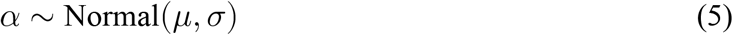

where *x*_*i*_ is the *i*th individual body size, *f*(*x*; λ, *x*_*min*_, *x*_*max*_, *counts*) is the truncated Pareto distribution, λ is the size spectrum parameter (also referred to as the exponent), *x*_*min*_, *x*_*max*_, and *counts* are as defined above, and *α* is the intercept with a prior probability distribution. In this case, we specified a Normal prior since λ is continuous and can be positive or negative.

For cases where the goal is to estimate changes in λ across space or time, the simple model above can be expanded to include predictors and/or varying intercepts and slopes:

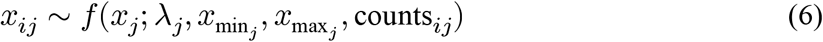

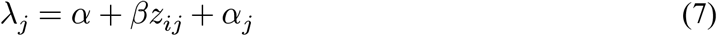

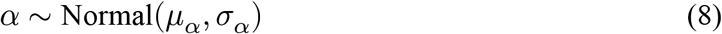

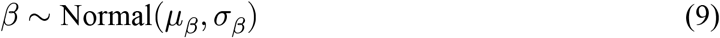

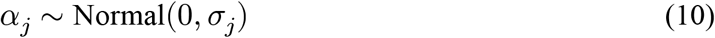

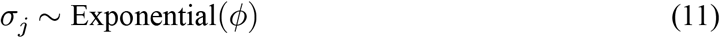

where *x*_*ij*_ is the *i*^th^ body size from group *j*. The groups might represent *j* sites, *j* experimental units, or *j* times, etc. The *x*_*ij*_ body sizes are distributed as a truncated Pareto with an unknown λ_*j*_, corresponding to the size spectrum parameter for each *j* group, along with group specific 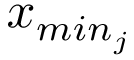and 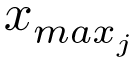 . The linear model for λ_*j*_ contains an intercept *α*, a slope *β*, a continuous predictor *z*_*ij*_, and a varying intercept for each group *α*_*j*_. In this example, prior distributions are *Normal* for *α, β*, and *α*_*j*_. *α* and *β* require priors for their respective mean *μ* and standard deviation *σ*. The varying intercept *α*_*j*_ has a mean = 0, and a standard deviation *σ*_*j*_ with it’s own *Exponential* hyperprior *ϕ*. The literature on prior choice is broad and active (Banner *et al*. 2020; Wesner & Pomeranz 2021), particularly for priors on hyperparameters like *σ*_*j*_ (Gelman 2006; Aguilar & Bürkner 2023). We specify prior distributions here for clarity, but users should choose prior distributions that reflect prior knowledge. An example of checking priors with the prior predictive distribution and prior sensitivity is in the Appendix (Figures S1 and S2).

### Testing the models

#### Parameter recovery from simulated data

To ensure that the models could recover known parameter values, we simulated *k* = 1000 data sets from a bounded power law using the inverse cumulative density function:

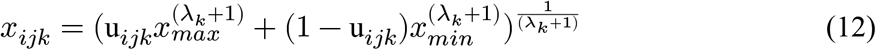

where *x*_*ijk*_ is the *i*^th^ individual body size from the *j*^th^ value of λ for the *k*^th^ simulation and j = 1,2,…,7 equally spaced lambda values from -2.4 to -1.2. u_*ijk*_ is a unique draw from a *Uniform*(0, 1) distribution, and all other variables are the same as defined above. We set *x*_*min*_ = 1, *x*_*max*_ = 1000, and simulated 300 body sizes (*i* = 1,2,3,…,300) for each *j* and *k*. To generate *counts*, we rounded each simulated value to the nearest 0.001 and tallied them.

##### Individual lambdas

After simulating the data, we estimated the λ values in three ways. First, we fit separate intercept-only models (Eq. 3) to each simulated data set. This represents the common procedure of estimating λ’s independently before using them in later analyses (e.g., Arranz *et al*. (2019); Pomeranz *et al*. (2022)). Second, we fit a single fixed effects model of the form:

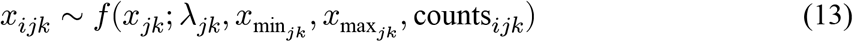

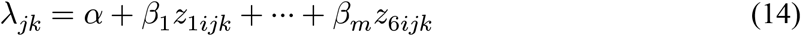

where *α* is the intercept representing the reference value of λ (in this case it is -2.4), *β*_*m*_ is the coefficient for the *m* = 1, 2, … 6 contrasts between the reference λ and λ_*m*_, and all other parameters and data are as described above. Third, we fit a varying intercepts model.

The procedures above resulted in 9000 total model fits (7000 for the separate λ estimates plus 1000 each for the fixed and varying intercept models). Each model fit includes newly simulated data from Eq. 13-14. To assess how well the models captured the known λ’s, we estimated coverage and bias for each λ. For coverage, we generated 95% credible intervals across each of the model runs and calculated the proportion of those CrI’s that contained the known λ value. For bias, we calculated the difference between the median estimated λ and the known λ across each model run.

##### Sample size and size range

We examined sensitivity to sample size (number of individual body sizes) across two λ values (-2, -1.6). For each λ, we simulated 30, 100, 300 or 1000 individuals. Each data set was fit using separate intercept-only models. We then repeated this process (data simulation and model fitting) n = 1000 times to estimate bias and coverage as described above.

In addition to sample size, we examined sensitivity to the size range, which can affect interpretations of λ (Sprules & Barth 2016). To do this, we again set λ to -2 or -1.6 and then simulated n = 300 individuals, varying *x*_*min*_ and *x*_*max*_ so that they contained 1, 2, 3, 4, or 5 orders of magnitude in range (i.e., *x*_*min*_ = 1 & *x*_*max*_ = 10/100/1000, etc.). For each size range, we repeated the data simulation and model fitting 1000 times to estimate bias and coverage. We also estimated precison as the range between the lowest and highest λ estimates across each simulation.

##### Regression

We simulated a linear regression model with a single continuous predictor *z* such that

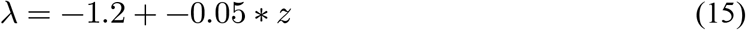

λ = -1.2 + -0.05*z*. This contains a known intercept *α* = -1.2 and a slope *β* = -0.05. The predictor *z* ranged from -1 to 1, with ten equally spaced intervals. Values for *α* and *β* were chosen to keep λ within typical ranges of -1 to -2 across the predictors. After obtaining the ten λ values (one for each value of *z*), we simulated 300 individuals from each λ using Eq. 12 and setting *x*_*min*_ = 1 and *x*_*max*_ = 1000. The regression was fit using Eqs. 6-8, but with the varying intercept *α*_*j*_. We repeated this procedure (data simulation and model fitting) 1000 times (*see Model Fitting*) and checked for parameter recovery, bias, and coverage as described above.

##### Benefit of partial pooling and priors

Using hierarchical Bayesian models has the benefit of improving λ and regression parameter estimates with partial pooling and informative priors. These can be especially important when data from different times or places have different sample sizes. To demonstrate this, we modified the linear regression described above to include 12 values of *z*, one of which was an “outlier” in which λ = -1.1 when *z* = 2.5. According to the regression equation, λ should actually equal -2.5 when *z* = 2.5. After estimating the λ’s, we again simulated n = 300 individuals from each lambda with *x*_*min*_ = 1 and *x*_*max*_ = 1000. However, for the outlier, we limited the number of individuals to n = 50. This mimics a situation in which an outlier is potentially due to a low sample size, a scenario for which partial pooling can be particularly effective (McElreath 2020 p. 413). The purpose of this exercise is not to reflect any particular sampling scheme, but to demonstrate the importance of partial pooling and priors.

We used three techniques to estimate the relationship between *z* and λ. First, we used the two-step process to 1) individually estimate each lambda, and 2) fit a Gaussian Bayesian linear regression between the between *z* and the separately estimated λ’s. This is akin to a no-pooling regression, in which no information (including about the sample size) is accounted for in the λ estimates. Second, we fit a linear mixed model with varying intercepts as described above, but with weak priors (*α ∼ N*(−1.5, 1), *β ∼ N*(0, 0.5), *σ*_*j*_ *∼ Exponential*(1). This model demonstrates partial pooling, in which the individual lambda estimates should be pooled towards the mean, particularly for the sample with 50 individuals. Additionally, the regression parameters (*α, β*), should be less influenced by the outlier compared to the first model. Finally, we fit a model with both varying intercepts and strong priors (*α ∼ N*(−1.5, 0.1), *β ∼ N*(−1, 0.02), *σ*_*j*_ *∼ Exponential*(1)).

### Model Fitting

Because the truncated Pareto pdf as described here is not available in rstan, we built an R package, isdbayes(Wesner & Pomeranz 2023), to integrate it into rstan using brms in R (Bürkner 2018; R Core Team 2020). The main benefit of brms is that it fits Bayesian models in rstan using common R modeling syntax. For example, this linear regression in R, lm(y ∼ x, data = data) becomes Bayesian using brm(y ∼ x, data = data), where brm will translate the model to rstan for MCMC sampling. The dots “…” indicate additional model specifications for the likelihood, priors, iterations, chains, etc. A short tutorial on using isdbayes is available at https://github.com/jswesner/isdbayes.

We specified each of the above models in brms, with the truncated Pareto added from the isdbayes package. Posteriors were explored in rstan (Stan Development Team 2022) using 2 chains each with 1000 iterations. All models converged with *R*_*hat*_’s <1.01. Assessments of prior influence and model checking are demonstrated in Appendix (Figure S3). In particular, for model checking, we use simulations from the posterior predictive distributions. These simulations can check how well the model resembles the raw data. Strong deviations from raw data may indicate poor model specification or may indicate deviation from the assumption of the power law.

## Data Availability Statement

All data, R code, and Stan code are available at https://github.com/jswesner/stan_isd (to be permanently archived on acceptance).

## Results

### Individual lambdas

The three methods (separate models, fixed effects, and varying intercepts) recaptured the true λ values (Figure 1) with no apparent evidence of bias. For example, mean bias ranged from -0.01 to 0.008, but all standard deviations included zero (Table 1). Similarly, coverage ranged from 0.93 to 0.96 with an grand mean of 0.95, indicating similarity to the nominal coverage of 0.95 (Table 1).

**Table 1:** Table 1. Parameter recovery using three modeling approaches with the same data. First, separate models individually recapture known lambda values. Second, lambdas are estimated using a single fixed effects model. Third, lambdas are estimated hierarchically using a single varying intercepts models. Each model and data simulation procedure is repeated 1000 times. Coverage is estimated for 95% Credible Intervals. Bias represents the mean and standard deviation of bias across the 1000 replicates.

| True Lambda | Metric | Separate Models | Fixed Predictor | Varying Intercepts |
| --- | --- | --- | --- | --- |
| -2.4 | Coverage | 0.93 | 0.94 | 0.95 |
| -2.2 | Coverage | 0.95 | 0.95 | 0.96 |
| -2.0 | Coverage | 0.95 | 0.96 | 0.95 |
| -1.8 | Coverage | 0.94 | 0.95 | 0.95 |
| -1.6 | Coverage | 0.95 | 0.95 | 0.95 |
| -1.4 | Coverage | 0.94 | 0.94 | 0.94 |
| -1.2 | Coverage | 0.96 | 0.95 | 0.95 |
| -2.4 | Bias | -0.004 (0.08) | -0.005 (0.08) | 0.008 (0.08) |
| -2.2 | Bias | -0.016 (0.07) | -0.007 (0.07) | 0.003 (0.07) |
| -2.0 | Bias | -0.004 (0.06) | -0.005 (0.06) | -0.001 (0.06) |
| -1.8 | Bias | -0.004 (0.05) | -0.002 (0.05) | -0.005 (0.05) |
| -1.6 | Bias | -0.005 (0.04) | -0.004 (0.04) | -0.004 (0.04) |
| -1.4 | Bias | -0.001 (0.03) | -0.003 (0.04) | -0.005 (0.04) |
| -1.2 | Bias | -0.001 (0.03) | -0.002 (0.03) | -0.004 (0.03) |

**Figure 1.**
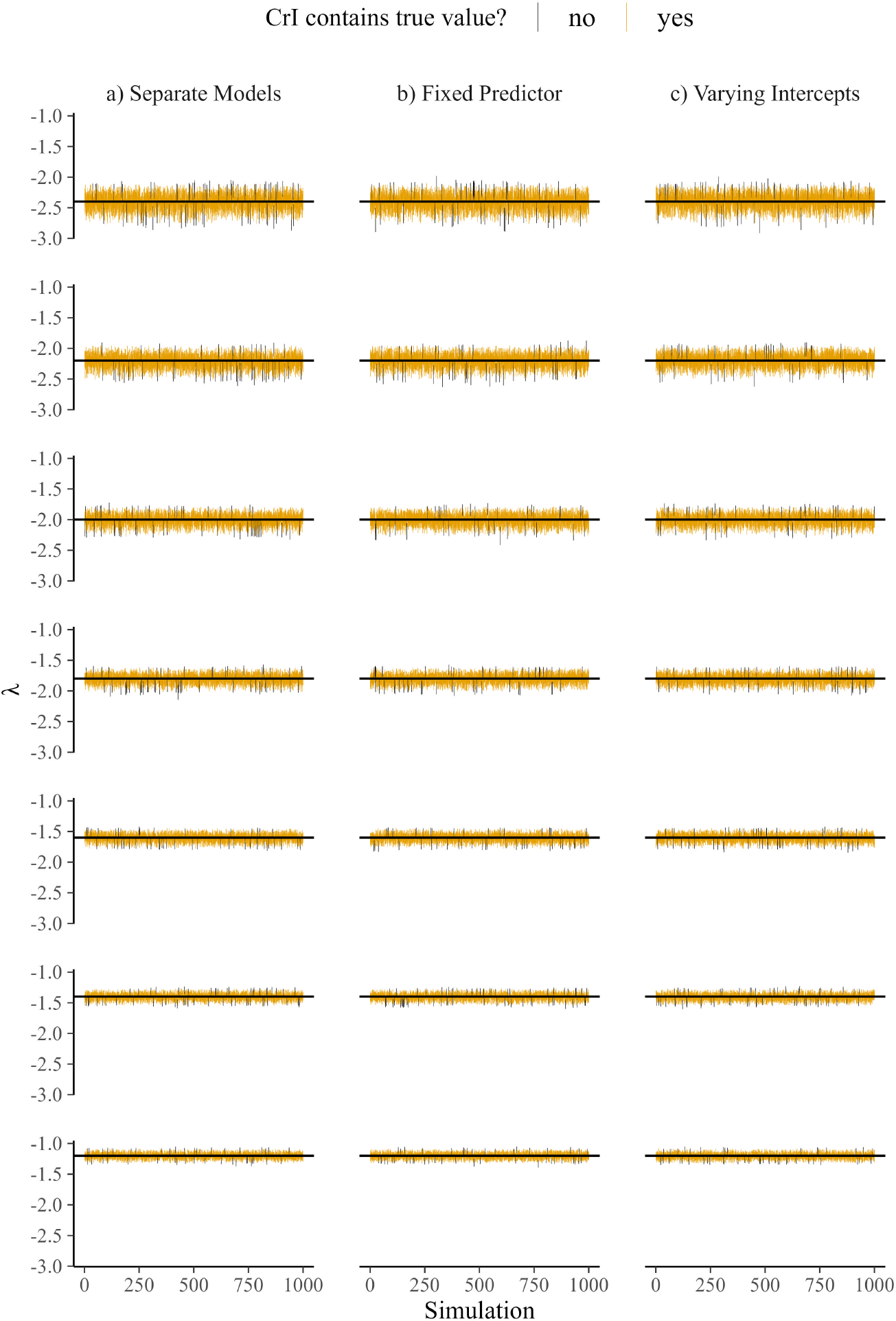
Modeled 95% credible intervals (CrI; n = 1000) of seven λs using a) separate interceptonly models for each lambda, b) a fixed linear predictor with the λ value as a group, and c) varying intercepts. Horizontal black lines show the true λ. Intervals either include the true λ (yellow) or not (black).

### Sample size and size range

Coverage ranged from 0.93 to 0.96 across sample sizes (Figure 2a),indicating good statistical coverage even at low sample sizes. However, precision increased with sample size. At n = 1000 and λ = -1.6, the range of mean λ estimates (largest minus smallest λ) was 0.13. By comparison, it was 0.9 at n = 30 (Figure 2a). In addition, at n = 30 there was a slight negative bias, with the mean λ 0.03 units smaller than the true λs on average (though the standard deviations all covered the true λ). This bias disappeared when when n >= 100 individuals (Figure 2a).

**Figure 2.**
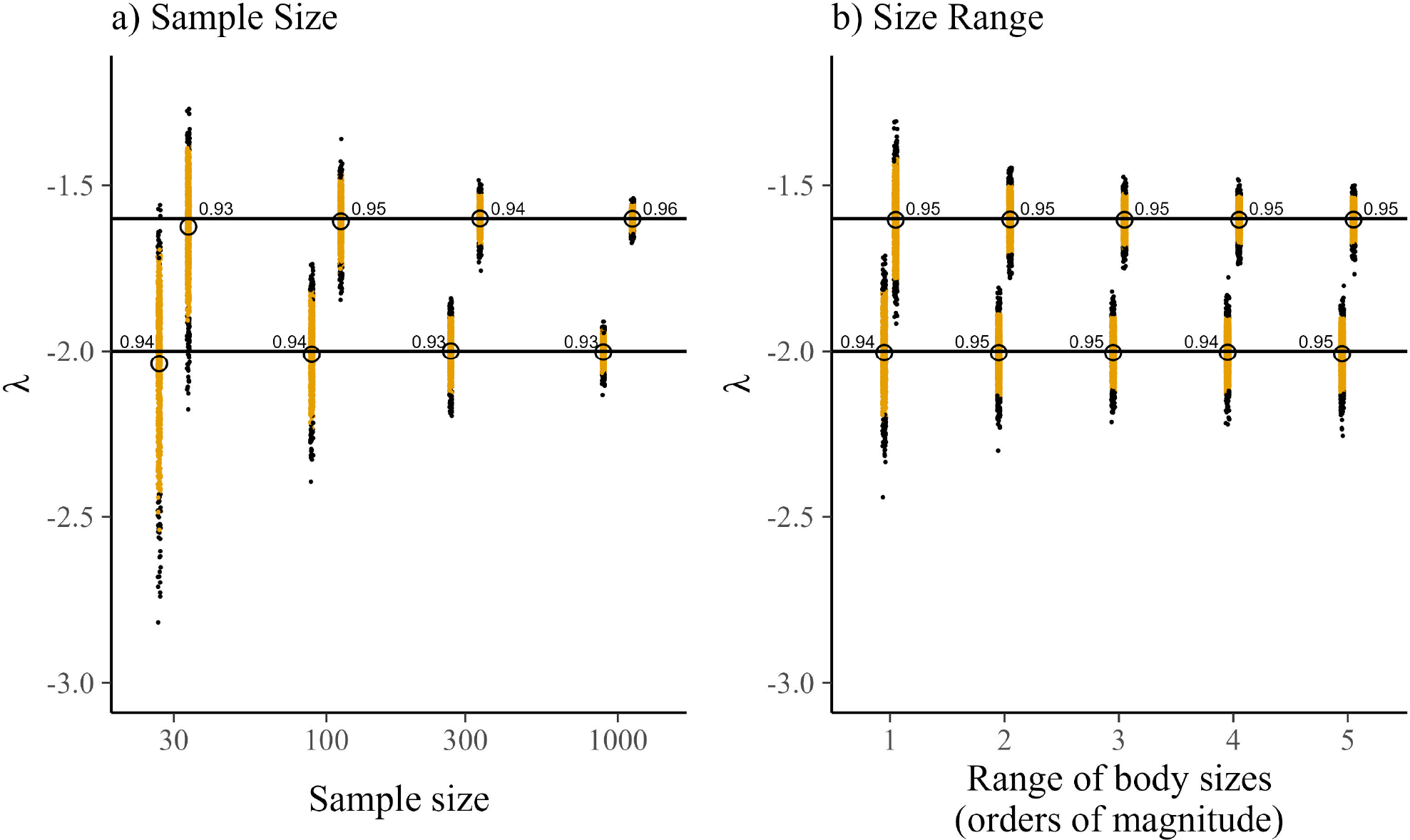
a) Changes in parameter estimation and coverage (numbers next to dots) as a function of sample size. Sample size is the number of individual body sizes used to estimate λ. Estimates of λ were repeated n = 1000 times for each sample size and known λ combination. b) The effect of size range on λ estimates. Modeled estimates (n = 1000) of two λs using separate intercept-only models with *x*_*min*_ and *x*_*max*_ ranging 1 to 5 orders of magnitude. Horizontal black lines show the true λs (-1.6 or -2). Closed dots in a) and b) are the posterior median λ estimates. 95% Credible intervals of those estimates (not shown for clarity) either include the true λ (yellow) or not (black). Open circles are the means for each group.

Coverage was also consistent across size ranges, achieving nominal coverage even at size ranges of 1 order of magnitude (Figure 2b). There was also no indication of bias, with mean λ estimates ranging from -0.002 to -0.007 units away from the true λ and standard deviations including λ. However, precision was lower when body sizes ranged 1 order of magnitude (range of estimates = 0.7 and 0.6 units for λ = -2 and -1.6, respectively). Precision declined to ∼ 0.4 and ∼ 0.3 at body size ranges of 2 or more orders of magnitude and remained relatively stable (Figure 2b).

### Regression

Coverage for the intercept (*α*) and slope (*β*) parameters was 95% (Table 2, Figure 3). Bias was small for both parameters, averaging -0.0003 for *α* and -3e-06 for *β*, indicating good parameter recovery.

**Table 2:**
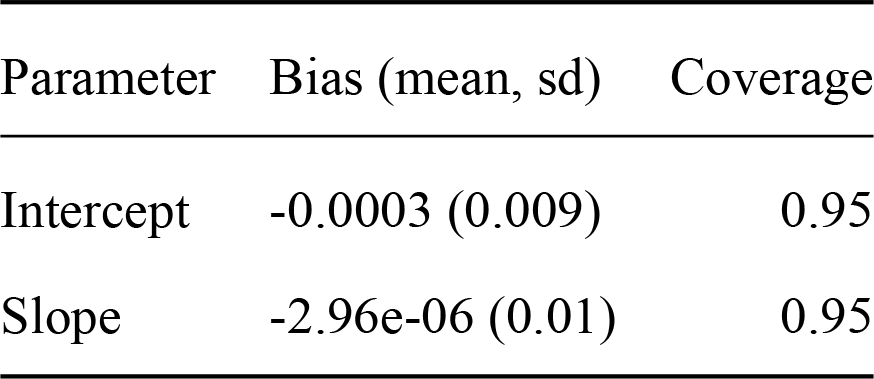
Bias and 95% coverage probabilities for the intercept and slope parametres of a linear regression. Value are estimated across 1000 simulations, where each simulation includes simulation of body sizes and a model fit using the isdbayes package.

| Parameter | Bias (mean, sd) | Coverage |
| --- | --- | --- |
| Intercept | -0.0003 (0.009) | 0.95 |
| Slope | -2.96e-06 (0.01) | 0.95 |

**Figure 3.**
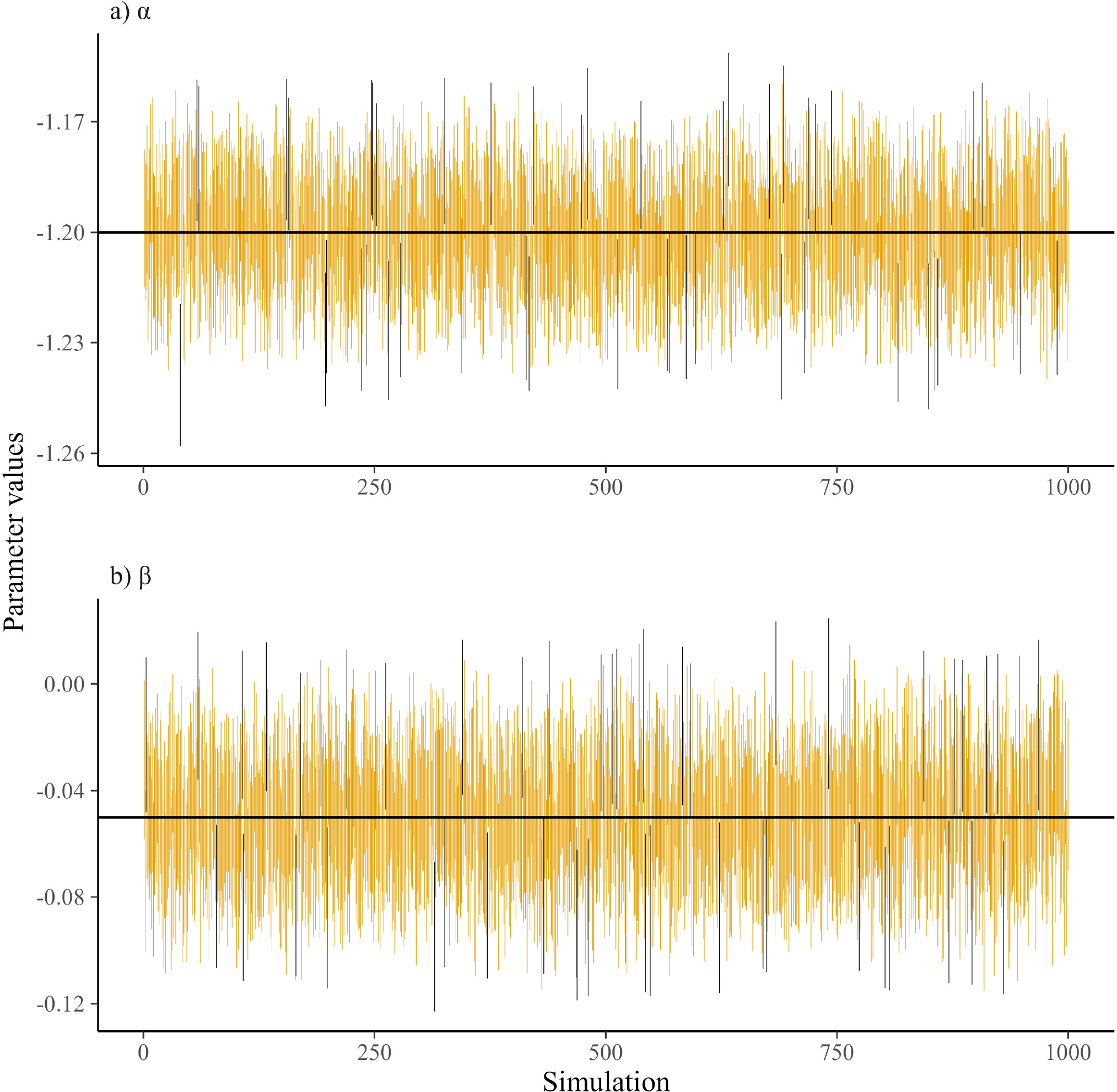
Modeled 95% credible intervals of the intercept (*α*) and slope (*β*) of a generalized linear regression estimating the change in λ across a predictor. Horizontal black lines show the true λ. Intervals either include the true λ (yellow) or not (black).

### Benefit of partial pooling and priors

Without partial pooling or informative priors, the two-step method was heavily influenced by the outlier, yielding a slope of -0.03 (95% CrI: -0.1 to 0.04; Figure 4a). While the credible interval contains the true slope (-0.1), there is high uncertainty in both the slope value and its sign. For example, there was only a 0.77 probability of a negative slope (and a 0.23 probability of a positive slope).

**Figure 4.**
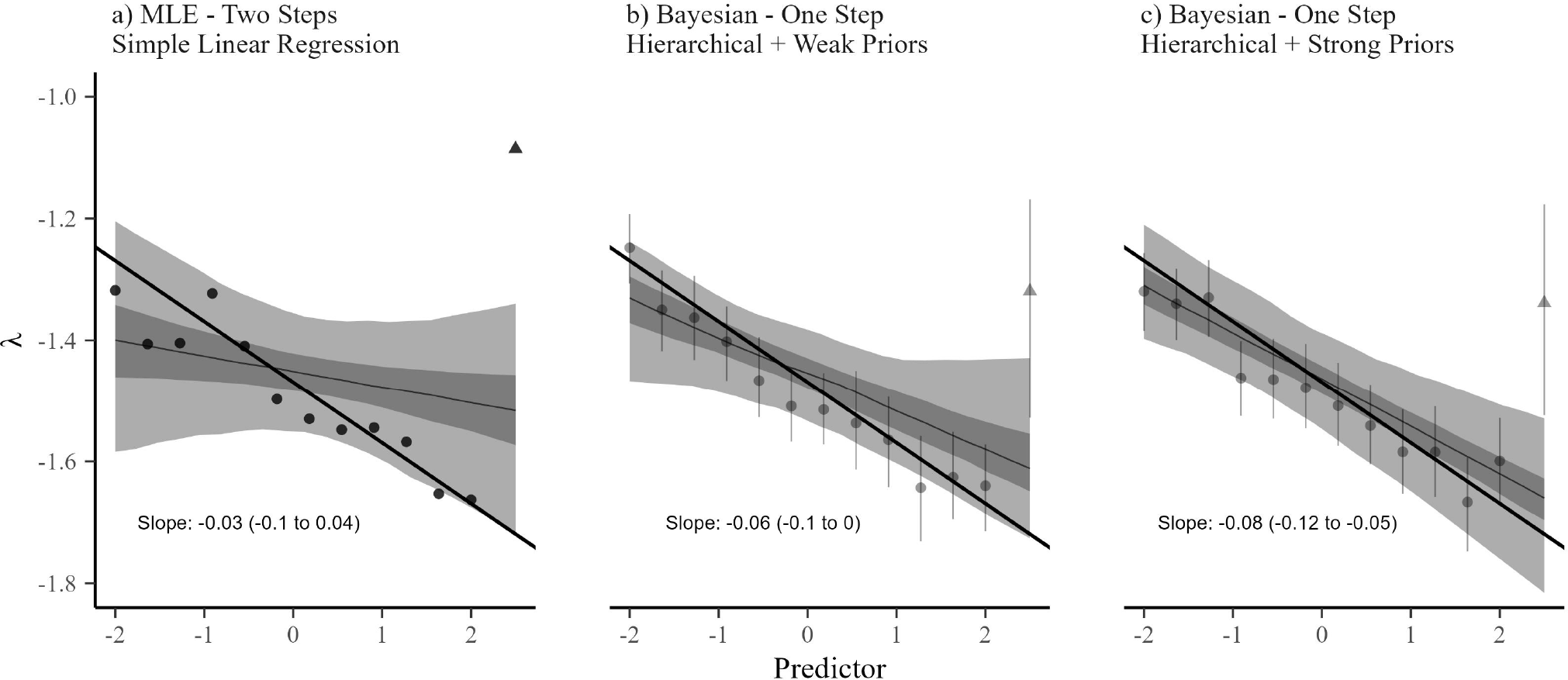
Regression results from a) a two-step process where λ’s are first estimated with separate models and then used as the response variable in a Gaussian regression, b) a generalized linear mixed model with a truncated Pareto likelihood and weak priors, and c) a generalized linear mixed model with a truncated Pareto likelihood and strong priors. The solid black line shows the true regression slope. Dark shading shows the 50% CrI and light shading shows the 95% CrI. All models have the same underlying individual body size data.

Fitting the same data with a single truncated Pareto linear mixed effects model reduced the influence of the outlier, yielding a slope of -0.06 (-0.1, to 0), 50% closer to the true slope of -0.1 than the two-step model (Figure 4b). In addition, this model more reliably captured the correct sign, with a 0.96 probability of a negative slope. Adding strong priors on the slope and intercept parameters further improved the estimate (Figure 4c), with a 0.99 probability of a negative slope.

In addition to improving parameter estimates, the λ estimates themselves are improved in the partially pooled models (Figure 4b,c). For example, in the two-step method, λ in the outlier is estimated -1.1 (Figure 4a), but it is reduced to ∼ -1.34 with partial pooling (Figure 4b,c). Partial pooling has a minimal effect on the other λ’s due to their larger underlying sample size.

## Discussion

The most important result of this work is the ability to analyze individual size distributions (ISD’s) using fixed and random predictors in a hierarchical model. Our approach allows ecologists to test hypotheses about size spectra while avoiding the pitfalls of a two-step process in which λ is estimated individually for each sample and the results are then used as response variables in linear or non-linear models. The generalized mixed model with a bounded truncated Pareto merges these steps, linking the data generation process (e.g., individual body sizes) with the model predictors. This permits the use of prior probabilities and hierarchical structure on regressions of ISD’s in a single analytical framework.

The ability to incorporate prior information using Bayesian updating has two practical advantages over the two-step process described above. First, adding informative prior distributions can improve model fit by limiting the MCMC sampler to reasonable sampling space. In other words it would not be sensible to estimate the probability that λ is -1,234 or -9. Without informative priors, those values (and more extreme values) are considered equally likely and hence waste much of the algorithm’s sampling effort on unlikely values (e.g., (Wesner & Pomeranz 2021)).

Second, and most importantly, ecologists have much prior information on the values that λ can take. For example, global analysis of phytoplankton reveals values of -1.75, consistent with predictions based on sub-linear scaling of metabolic rate with mass of -3/4 (Perkins *et al*. 2019). Alternatively, Sheldon’s conjecture suggests that λ is -2.05 (Andersen et al. 2006), a value reflecting isometric scaling of metabolic rate and mass, with support in pelagic marine food webs (Andersen & Beyer 2006). However, benthic marine systems typically have shallower exponents (e.g., *∼* -1.4; Blanchard *et al*. (2009)), similar to those in some freshwater stream ecosystems (*∼* -1.25(Pomeranz *et al*. 2022). While the causes of these deviations from theoretical predictions are debated, it is clear that values of λ are restricted to a relatively narrow range between about -2.05 and -1.2. But this restriction is not known to the truncated Pareto, which has no natural lower or upper bounds on λ (White *et al*. 2008). As a result, a prior that places most of its probability mass on these values (e.g., *Normal*(−1.75, 0.2) seems appropriate. Such a continuous prior does not prevent findings of larger or smaller λ, but instead places properly weighted skepticism on such values.

An important assumption when setting priors is that we have a good understanding of the values that λ can reasonably take. For most of the examples here, our priors are weakly informative in the sense that they rule out clearly unreasonable values (e.g., λ = -25, etc), but have weak effects on values within reasonable ranges (e.g., λ *∼* -3 to 0). Most published values of λ fall into this range regardless of the method used by those studies to estimate λ (White *et al*. 2007; Edwards *et al*. 2017). However, if more informative priors are required, such as our example in Figure 4c, then caution should be used when comparing prior expectations to previously estimated λs. For example, in an analysis of marine fish trawl data, Edwards *et al*. (2020) found that binning methods produced λ estimates of ∼ -2.2 across 30 years of data. Yet re-analysis of the same data using the truncated Pareto found λ estimates closer to -1.6. If we were to use these values to guide prior selection, then the choice of reasonable prior would clearly depend on the method used to estimate λ. The simplest approach would be to assume a fixed correction between the binning methods and the truncated Pareto when setting priors based on binning methods. Unfortunately, such a fixed correction does not appear to exist (Pomeranz *et al*. 2023), making it difficult or impossible to use λs from binning methods to guide informative prior selection.

Similar to priors, partial pooling from varying intercepts provides additional benefits, allowing for the incorporation of hierarchical structure and pulling λ estimates towards the global mean (Gelman 2005; Qian *et al*. 2010). In the example shown here, pooling was able to downweight the influence of an outlier that had a relatively small sample size (n = 50 individuals compared to n = 300). By contrast, in the two step-method, the same outlier had a large influence on the regression outcome, because the model had no information on the number of individuals used to generate each λ. Another benefit of pooling (both from varying effects and skeptical priors) is in prediction (Gelman 2005; Hobbs & Hooten 2015). This becomes especially important when models are used to forecast future ecosystem conditions. Forecasts are becoming more common in ecology (Dietze *et al*. 2018) and are likely to be easier to test with modern long-term data sets like NEON (National Ecological Observatory Network) in which body size samples will be collected at the continental scale over at least the next 20 years (Kuhlman *et al*. 2016). In addition, because the effects of priors and pooling increase with smaller samples sizes, varying intercepts are likely to be particularly helpful for small samples. In other words, priors and partial pooling contain built-in skepticism of extreme values, ensuring the maxim that “extraordinary claims require extraordinary evidence”.

One major drawback to the Bayesian modeling framework here is time. Bayesian models of even minimal complexity must be estimated with Markov Chain Monte Carlo techniques. In this study, we used the No U-Turn sampling (NUTS) algorithm (Hoffman *et al*. 2014) via rstan (Stan Development Team 2022). Stan can be substantially faster than other commonly used programs such as JAGS and WinBUGS, which rely on Gibbs sampling. For example, Stan is 10 to 1000 times more efficient than JAGS or WinBUGS, with the differences becoming greater as model complexity increases (Monnahan *et al*. 2017). In the current study, intercept-only models for individual samples with *∼* 300 to 1500 individuals could be fit quickly (<2 seconds total run time (warm-up + sampling on a Lenovo T490 with 16GB RAM)) with as little as 1000 iterations and two chains. However, the hierarchical regression models took >1 hour to run with the same iterations and chains. These times include the fact that our models used several optimization techniques, such as weakly informative priors, standardized predictors, and non-centered parameterization, each of which are known to improve convergence and reduce sampling time (McElreath 2020). But if Bayesian inference is desired, these run-times may be worth the wait. In addition, they are certain to become faster with the refinement of existing algorithms and the introduction of newer ones like Microcanonical HMC (Robnik *et al*. 2022).

Body size distributions in ecosystems have been studied for decades, yet comprehensive analytical approaches to testing these hypotheses are lacking. We present a single analytical approach that takes advantage of the underlying data structures of individual body sizes (Pareto distributions) while placing them in a generalized (non)-linear hierarchical modeling framework. In addition to fitting regression models, the results suggests that sample sizes >300 individuals, but optimally >1000, are sufficient to accurately estimate λ. We also found good performance at size ranges from 2 to 5 orders of magnitude, though it is important that this result is based on simulated data in which *x*_*max*_ is known and sampled with high probability. In a field sample that ranges, say, 3 orders of magnitude in body size, researchers should ensure that this range reflects the likely range of true sizes in the data set. We hope that ecologists will adopt and improve on the models here to critically examine hypotheses of size spectra or other power-law distributed data. Moreover, while the examples here are for ecological size spectra, the statistical approach is not limited to ecological data, but can be applied to analysis of power law distributions that are common in a wide variety of disciplines (Aban *et al*. 2006; Clauset *et al*. 2009).

## Acknowledgements

This material is based upon work supported by the National Science Foundation under Grant Nos. 2106067 to JSW and 2106068 to JRJ. We thank Professor Yuhlong Lio for statistical and mathematical advice and Edwards et al. (2017) and (2020) for placing their code and data in easily accessible repositories.

## Translating to Stan and brms

We translated the probability density function described by Edwards *et al*. (2020) into Stan by converting it to the log probability density function (lpdf). Stan is a probabilistic modeling language that is capable of fitting complex models, including those with custom lpdf’s. The resulting lpdf is given as

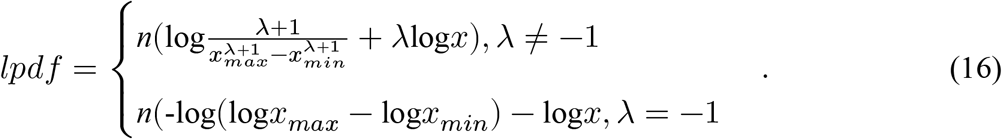

where all variables are as described in the main text. To make it easier to use, we also wrote the above *lpdf* as an custom response distribution for the R package brms. It is available at https://github.com/jswesner/isdbayes. For example, after cloning the repository for *isdbayes*, a model with a single predictor *group* can be coded in brms as follows:

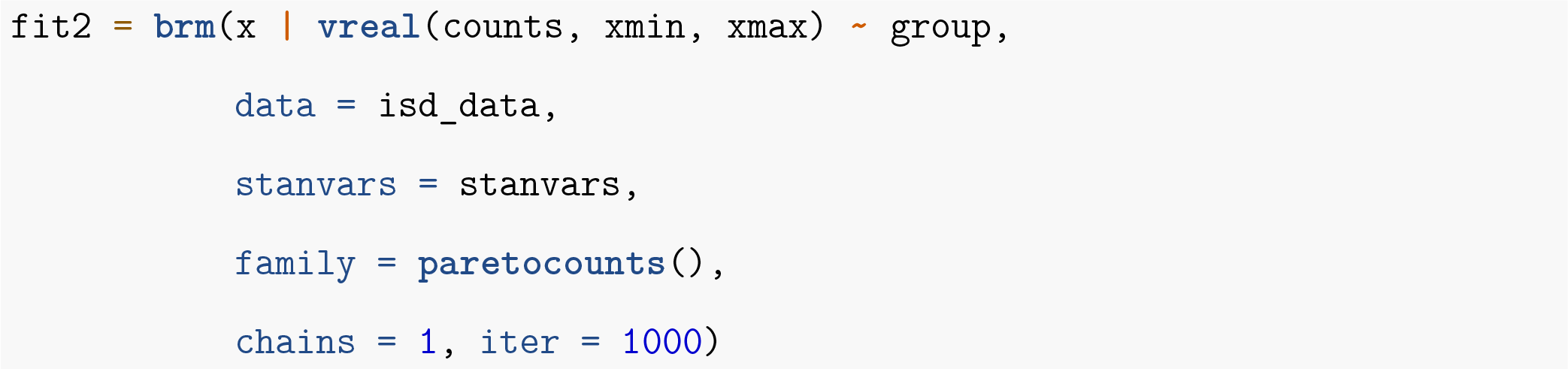

In the code above, the model equation on the right hand side of *∼* can take any number of forms that are consistent with standard model coding in R, including the addition of varying intercepts and varying slopes.

### Appendix S2

#### Model Assessment

Model assessment is a key part of the Bayesian workflow, one goal of which is to assess prior influence and model fit Gelman *et al*. (2020). Here, we provide examples of model assessment using the prior predictive distribution, a prior sensitivity analysis, and the posterior predictive distribution (Gabry *et al*. 2019). As model assessment is a broad and developing area (Conn *et al*. 2018), we do not intend this demonstration to be exhaustive, but merely to demonstrate it for the models developed in the main manuscript.

#### Prior Sensitivity

Prior implications are often best understood through visualizations of the prior predictive distribution Wesner & Pomeranz (2021). We demonstrate that here by simulating 300 body sizes from λ = -1.6. We then fit these data with an intercept only model using a Normal prior that varied in both mean (-2, -1.8, -1.5, -1.2) and standard deviation (10, 5, 2, 1, 0.1, 0.01, 0.001). The smaller standard deviations indicate increasingly informative priors. For each combination of priors, we simulated data and fit the model using the isdbayes package and brms. The result is shown in Figure S1. For highly informative prior values of 0.001 and 0.01, there is little influence of the data (Figure S1). However, for all combinations of the prior mean, the data are less influential when the prior sd is 0.1 or larger. In other words, a prior of ∼ 0.1 or larger has minimal influence on the λ estimates.

#### Prior Predictive

Figure S2 shows an example of visualizing the influence of the prior on the regression described in Figure 4b in the main text. In this case, the priors for the regression were Normal(-1.5, 1) for the intercept and Normal(0, 0.5) for the slope. The model also has an exponential prior of exp(1) for the hyperprior standard deviation, but we excluded it here for clarity. Simulating 100 regression lines from the prior and posterior demonstrates the effect of the data, showing far less spread in possible regression lines in the posterior compared to the prior (Figure S2).

#### Model Fit

Because Bayesian models are generative, we can assess model fit by simulating data from the posterior distribution and comparing it to the raw data (Gabry *et al*. 2019). The motivation for this approach is that a model that faithfully recaptures the data generation process should generate data that resemble the raw data in at least some aspects. We did this by first fitting ISD’s to data generated from three known λ’s (-2, -1.6, -1.3). Then we used the posterior predictive distribution to simulate 500 datasets and compare them to the raw data. We did this visually using boxplots (Figure S3a-c) of the raw data compared to the first 10 simulations of data from the posterior. In addition, we calculated the geometric mean (GM) of each simulation and the raw data using

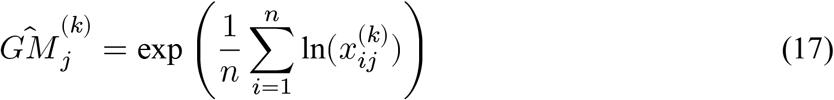

Where *x*_*ij*_ is the *i*^th^ body size from the *j*^th^ group, represented here by the different known lambdas used to simulate data (-2, -1.6 or -1.3). We generated a unique geometric mean for each of 500 posterior draws (*k*). We visualized discrepancies from the raw geometric mean using histograms (Figure S3d-f).

The results indicate good fit, showing that the posteriors generated from models strongly resemble the raw data (Figure S3a-c). In addition, the geometric means of the raw data were captured by the geometric means simulated from the posteriors (Figure S3d-f). One possible discrepancy appears in Figure S3f, where the true geometric mean is in the upper tail of the posterior histogram. A likely explanation for this is that the prior l is set to *N* (-1.8, 2), while the true l in Figure S3f is -1.3. It is possible that the discrepancy is due to the prior influence, though this should be explored. The most important result of this is simply to demonstrate that standard model assessment applies to the power-law models presented here. For a more thorough treatment of model checking, see Gelman *et al*. (2013) and Conn *et al*. (2018).

**Figure 1.**
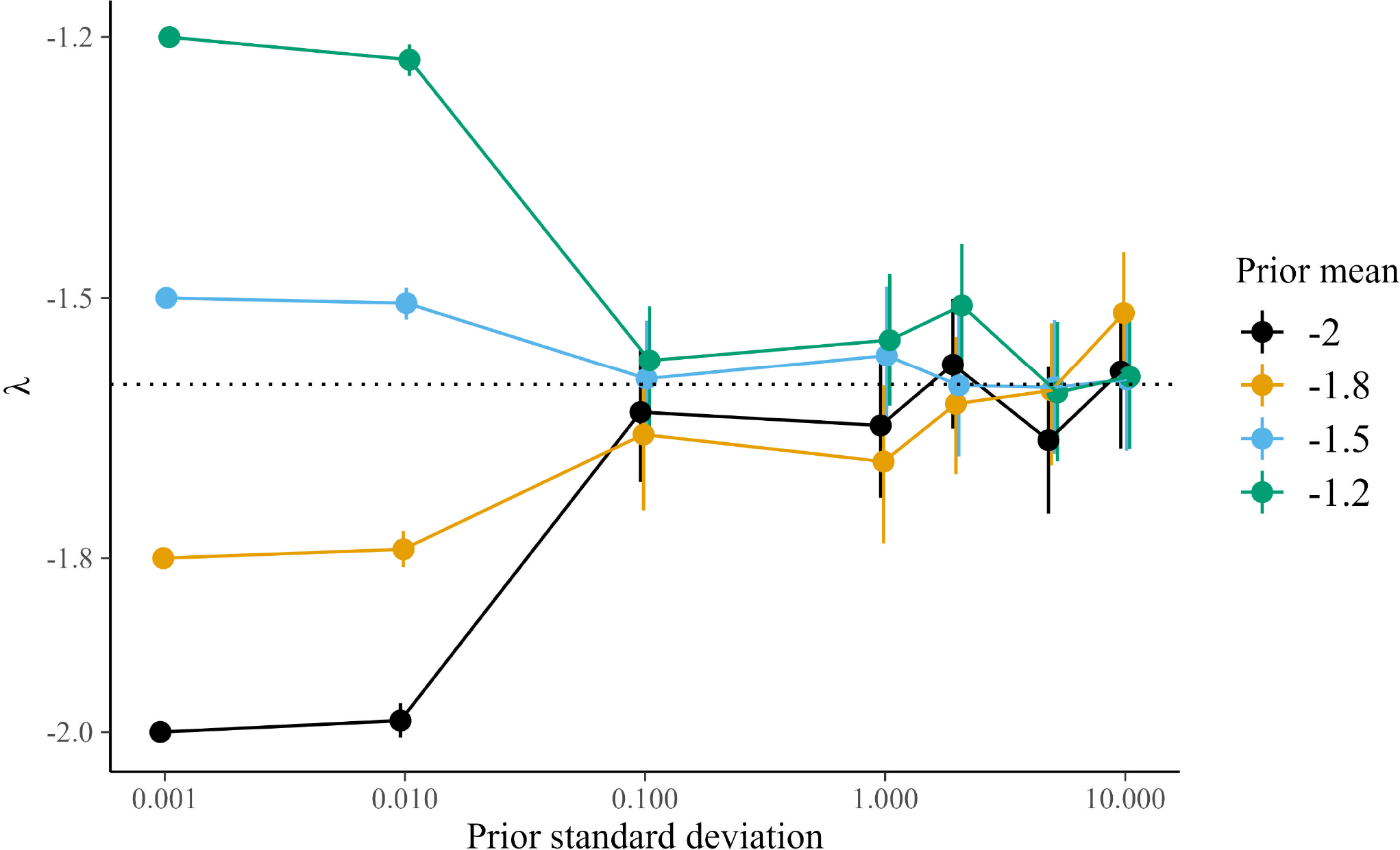
Prior sensitivity. The data are simulated from lambda = -1.6, shown by the dotted black line. Estimating those data from models with different prior means and standard deviations shows the influence of the standard deviation (x-axis).

**Figure 2.**
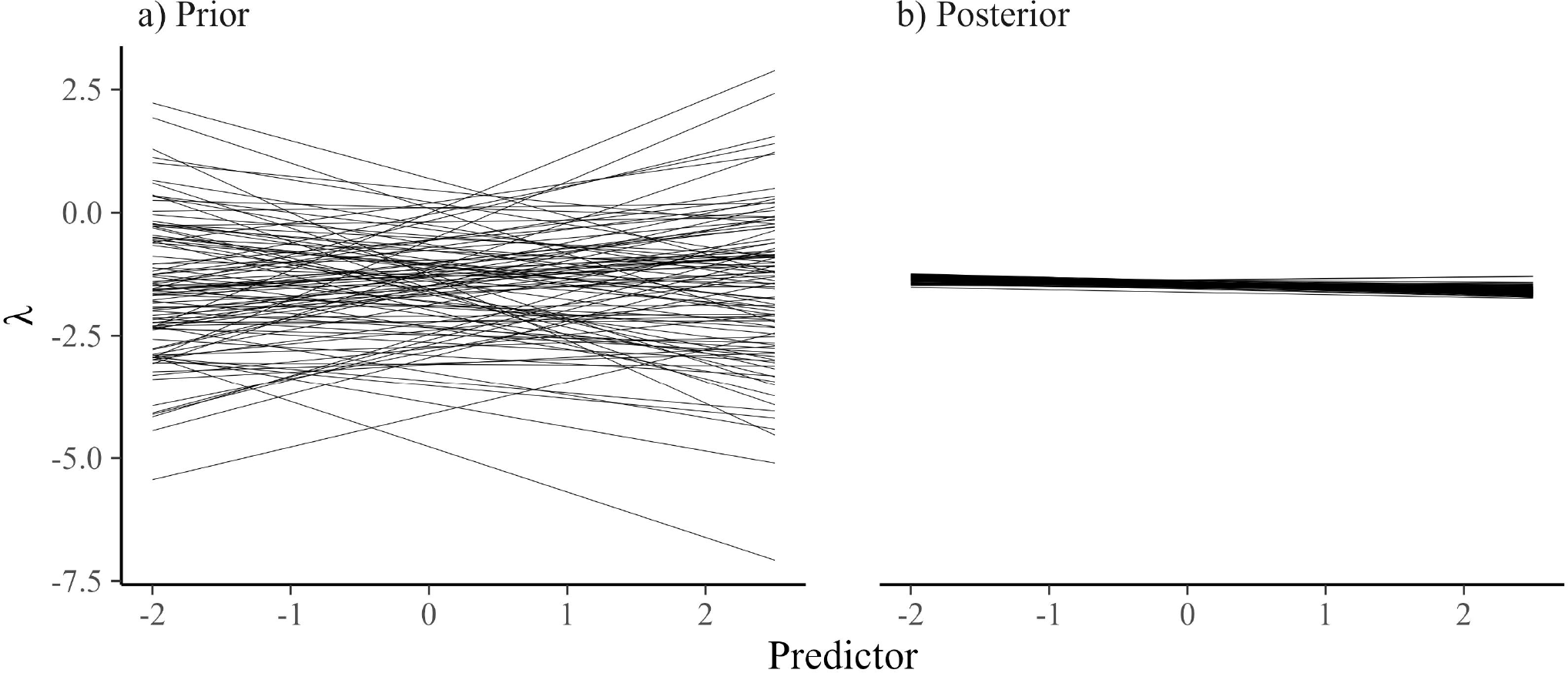
One hundred simulations from a) the prior distribution and b) the posterior distribution after fitting the model to data. Each line represents a single draw from the prior or posterior distributions.

**Figure 3.**
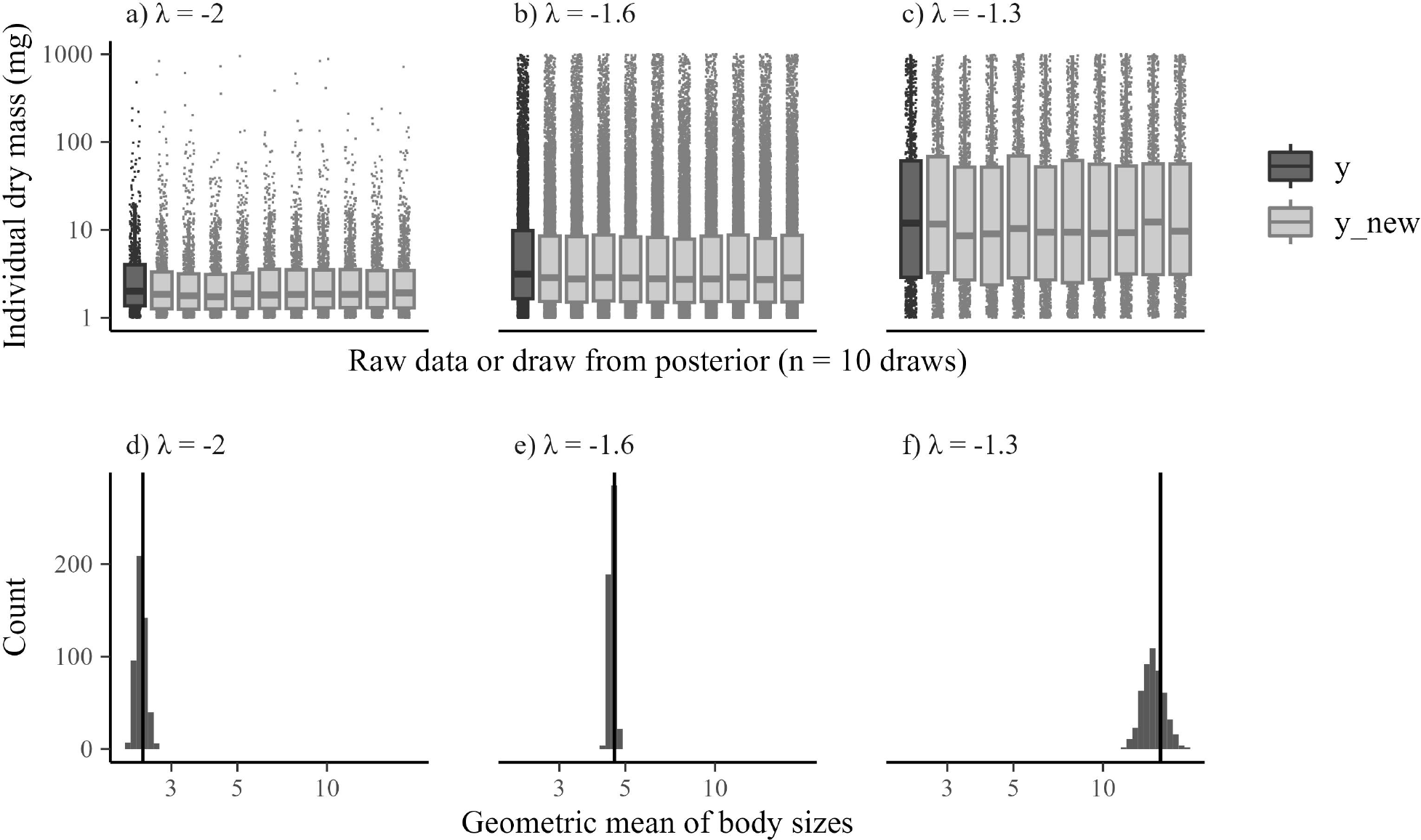
Posterior predictive checks of models estimating three ISDs with true lambdas ranging from -2 to -1.3. a-c) Raw data and boxplots from the original data (y) and 10 simulated datasets from the posterior *y*_*new*_. d-f) Histograms of the geometric mean from 500 simulated datasets from the posterior compared to the geometric mean of the raw data shown by the vertical line.

