## Appendix for "Bayesian hierarchical modeling of size spectra"

#### Appendix S1

*Translating to Stan and brms*

We translated the probability density function described by Edwards *et al.* (2020) into Stan by converting it to the log probability density function (lpdf). Stan is a probabilistic modeling language that is capable of fitting complex models, including those with custom lpdf's. The resulting lpdf is given as

$$lpdf = \begin{cases} n(\log\frac{\lambda+1}{x_{max}^{\lambda+1}-x_{min}^{\lambda+1}} + \lambda\log x), \lambda \neq -1 \\ n(\text{-}\log(\log x_{max} - \log x_{min}) - \log x, \lambda = -1 \end{cases} . \quad (1)$$

where all variables are as described in the main text. To make it easier to use, we also wrote the above *lpdf* as an custom response distribution for the R package `brms`. It is available at https://github.com/jswesner/isdbayes. For example, after cloning the repository for *isdbayes*, a model with a single predictor *group* can be coded in `brms` as follows:

```
fit2 = brm(x | vreal(counts, xmin, xmax) ~ group,
           data = isd_data,
           stanvars = stanvars,
```

```
            family = paretocounts(),
            chains = 1, iter = 1000)
```

In the code above, the model equation on the right hand side of $\sim$ can take any number of forms that are consistent with standard model coding in R, including the addition of varying intercepts and varying slopes.

$$\hat{GM}_j^{(k)} = \exp\left(\frac{1}{n}\sum_{i=1}^{n} \ln(x_{ij}^{(k)})\right) \tag{2}$$

Where $x_{ij}$ is the $i^{\text{th}}$ body size from the $j^{\text{th}}$ group, represented here by the different known lambdas used to simulate data (-2, -1.6 or -1.3). We generated a unique geometric mean for each of 500 posterior draws ($k$). We visualized discrepancies from the raw geometric mean using histograms (Figure S3d-f).

The results indicate good fit, showing that the posteriors generated from models strongly resemble the raw data (Figure S3a-c). In addition, the geometric means of the raw data were captured by the geometric means simulated from the posteriors (Figure S3d-f). One possible discrepancy appears in Figure S3f, where the true geometric mean is in the upper tail of the posterior histogram. A likely explanation for this is that the prior l is set to $N$(-1.8, 2), while the true l in Figure S3f is -1.3. It is possible that the discrepancy is due to the prior influence, though this should be explored. The most important result of this is simply to demonstrate that standard model assessment applies to the power-law models presented here. For a more thorough treatment of model checking, see Gelman *et al.* (2013) and Conn *et al.* (2018).

### Figures

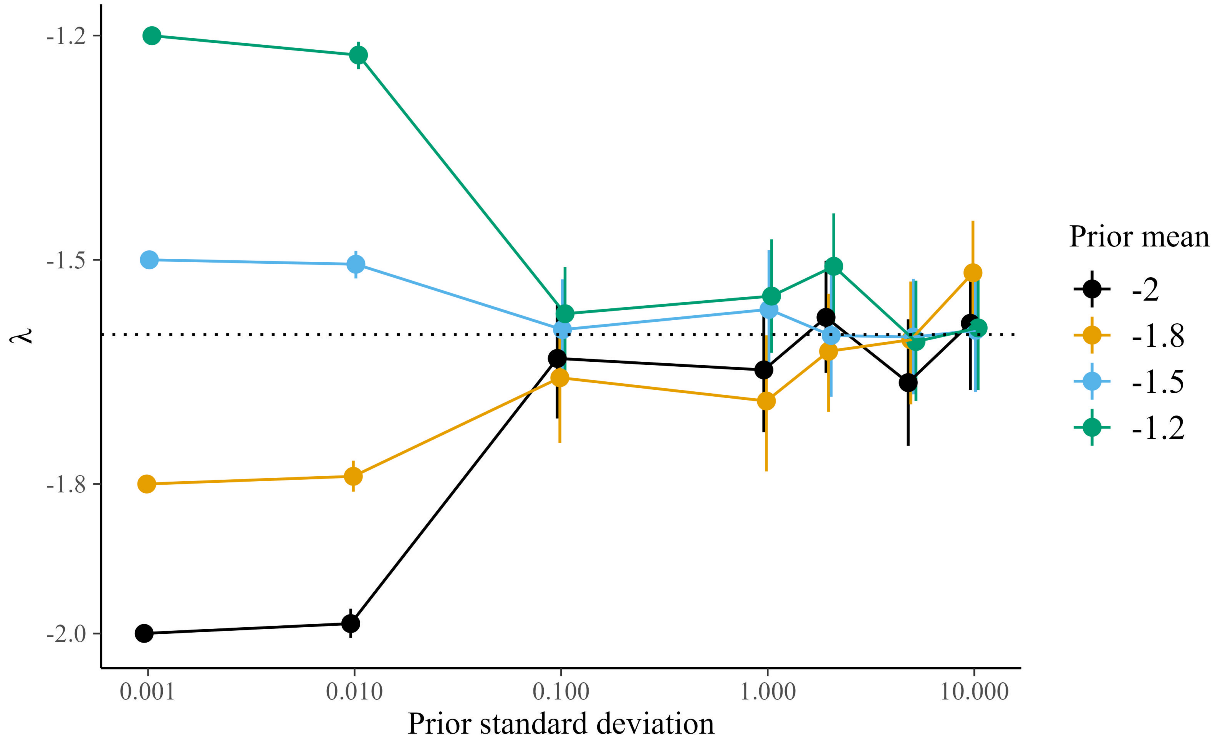


Figure S 1: Prior sensitivity. The data are simulated from lambda = -1.6, shown by the dotted black line. Estimating those data from models with different prior means and standard deviations shows the influence of the standard deviation (x-axis).

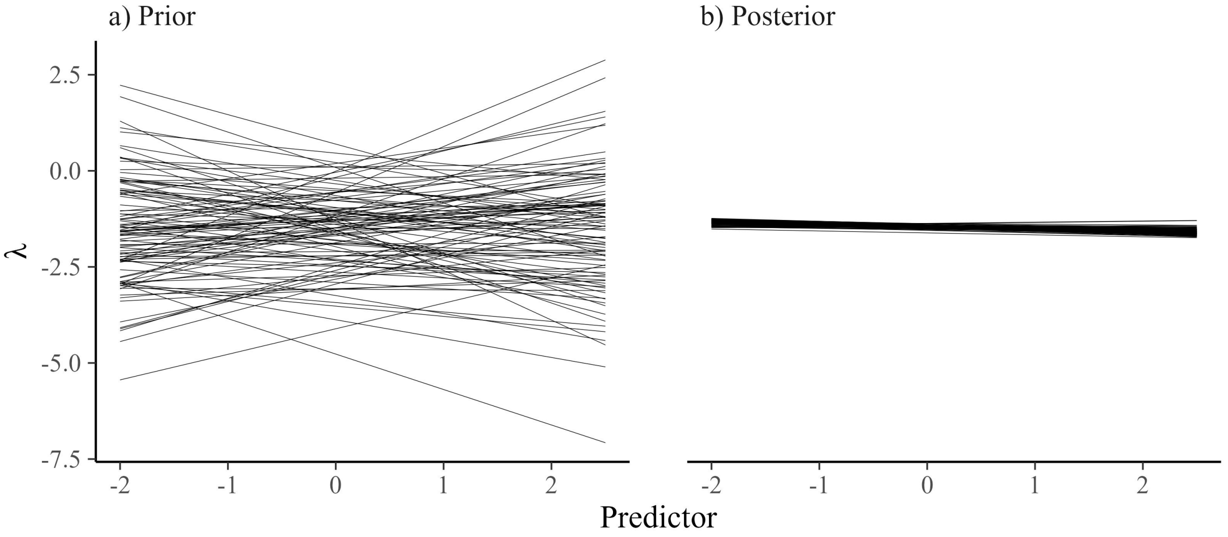


Figure S 2: One hundred simulations from a) the prior distribution and b) the posterior distribution after fitting the model to data. Each line represents a single draw from the prior or posterior distributions.

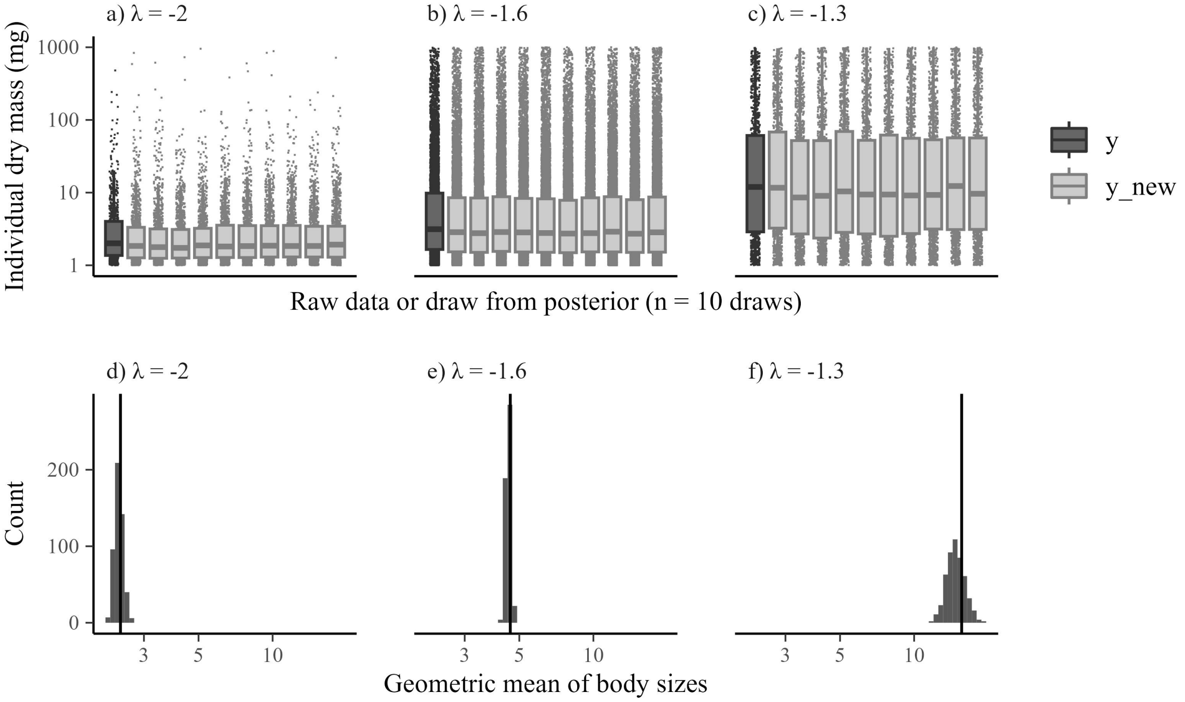


Figure S 3: Posterior predictive checks of models estimating three ISDs with true lambdas ranging from -2 to -1.3. a-c) Raw data and boxplots from the original data (y) and 10 simulated datasets from the posterior $y_{new}$. d-f) Histograms of the geometric mean from 500 simulated datasets from the posterior compared to the geometric mean of the raw data shown by the vertical line.
